## Supporting Information for "DeLTA: Automated cell segmentation, tracking, and lineage reconstruction using deep learning"

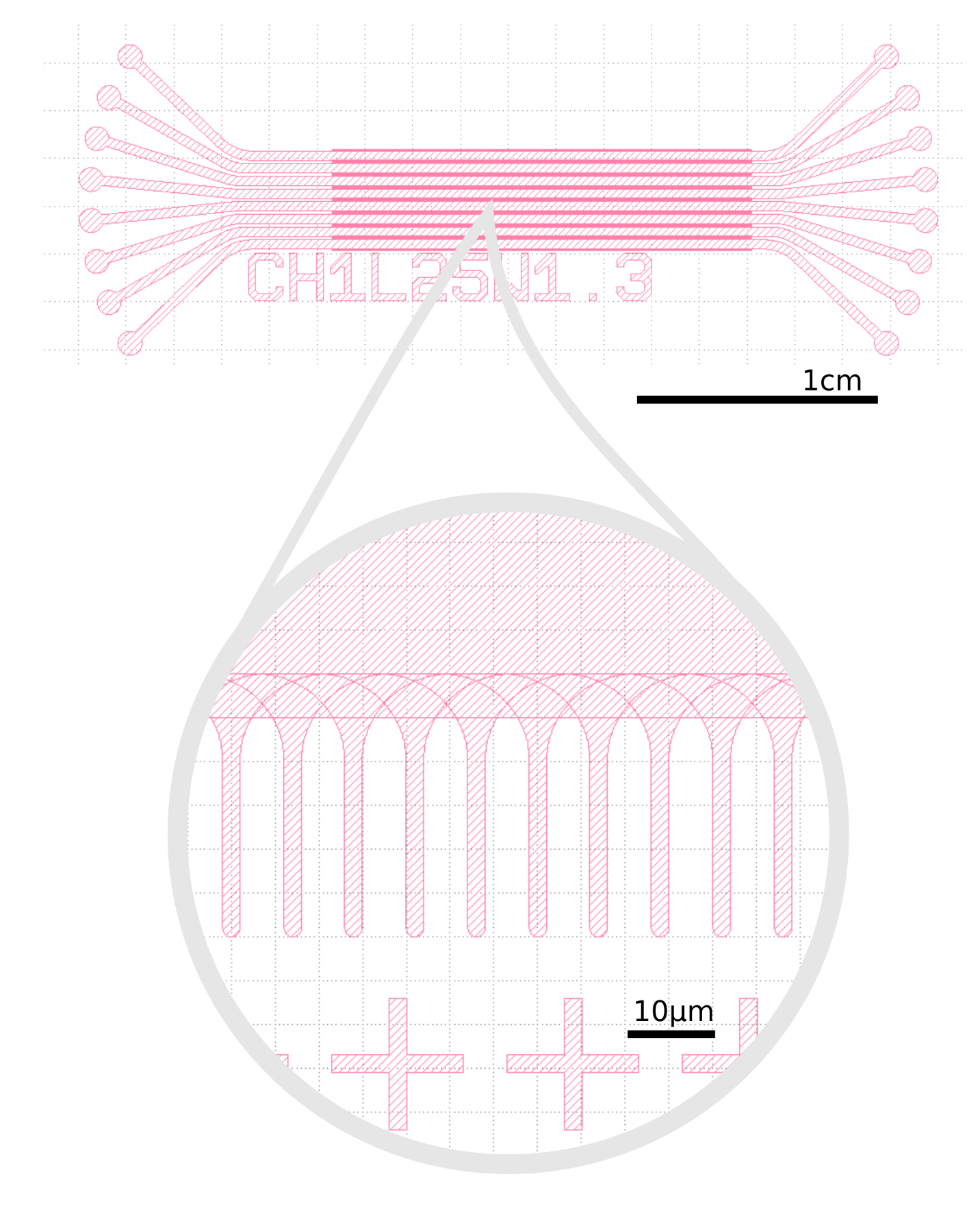

**Figure S1. Mother machine chip design used in this study.** The chip features 8 independent main channels of 400µm in width and ~60µm in height. Each channel features 6,000 chambers of 1.1µm in height, 25 or 35µm in length and 1.3 to 1.8µm in width.

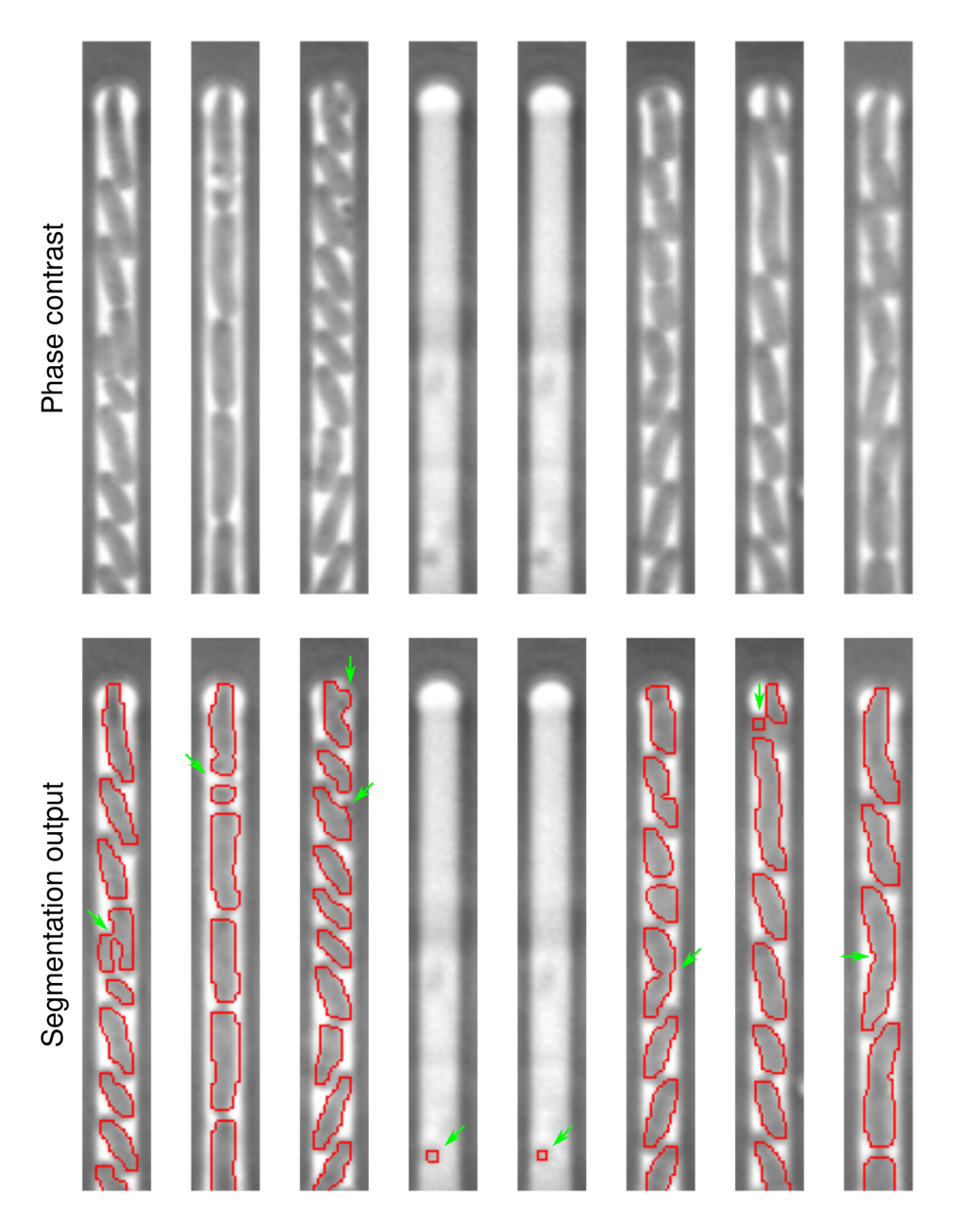

**Figure S2. Segmentation errors identified in our evaluation set.** 9 errors were identified out 6,311 segmented cells in the set. Errors are highlighted with green arrows.

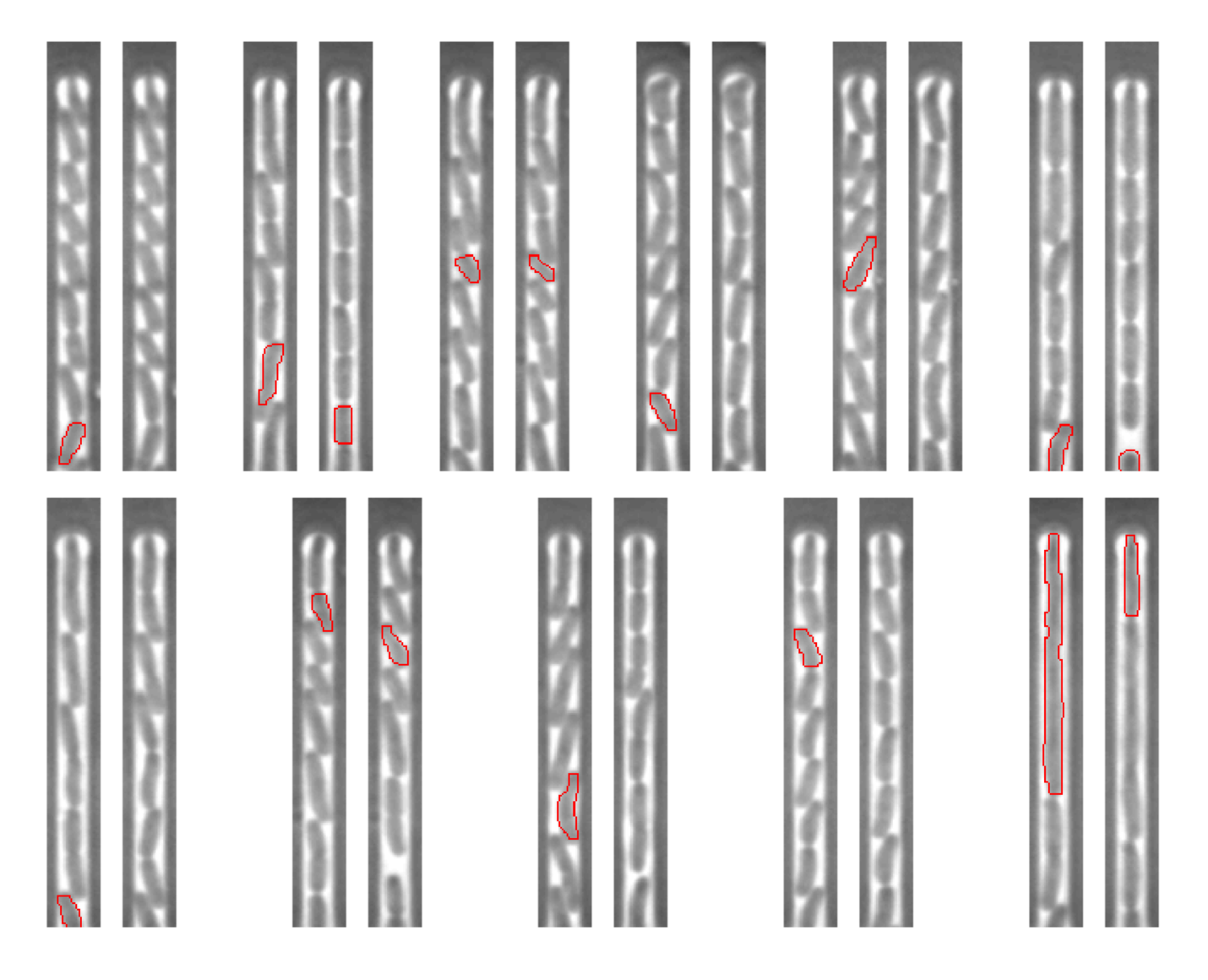

**Figure S3. Tracking errors identified in our evaluation set.** 11 errors were identified out of 1,040 tracking events in the set. Each image pair shows two subsequent time points. Red outlines show the tracked cells. Errors include cases where the algorithm fails to identify a cell in the subsequent frame, where the cell in the following frame is misidentified as part of the lineage, or where segmentation errors lead to tracking mistakes.

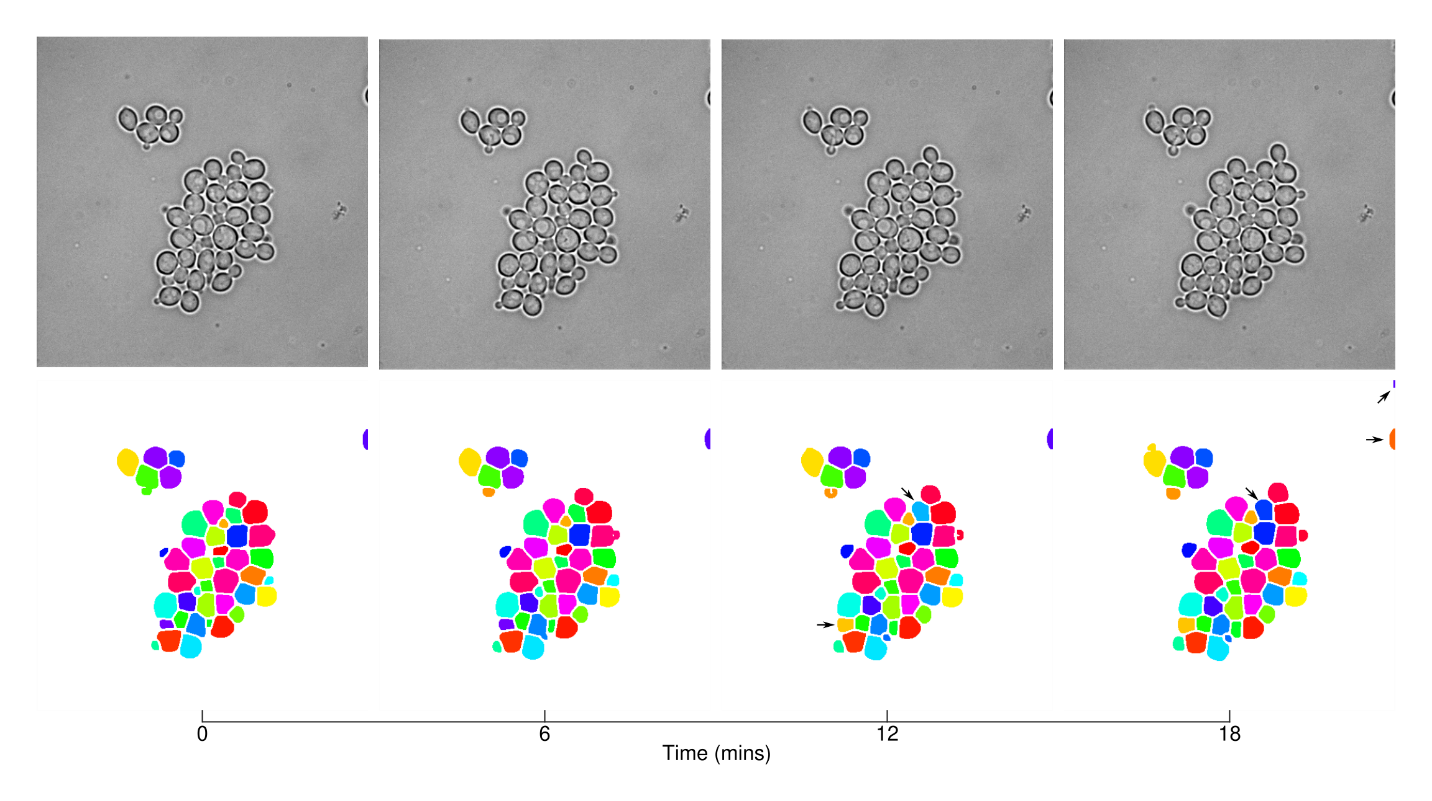

**Figure S4. Representative examples of yeast segmentation and tracking with DeLTA.** Arrows highlight tracking and segmentation errors. Note that the original training sets did not feature mother-daughter relationship information so we did not train the tracking U-Net to identify daughter cells. The two U-Net models were not trained on data from the experiment shown here, but on other experiments in the dataset.

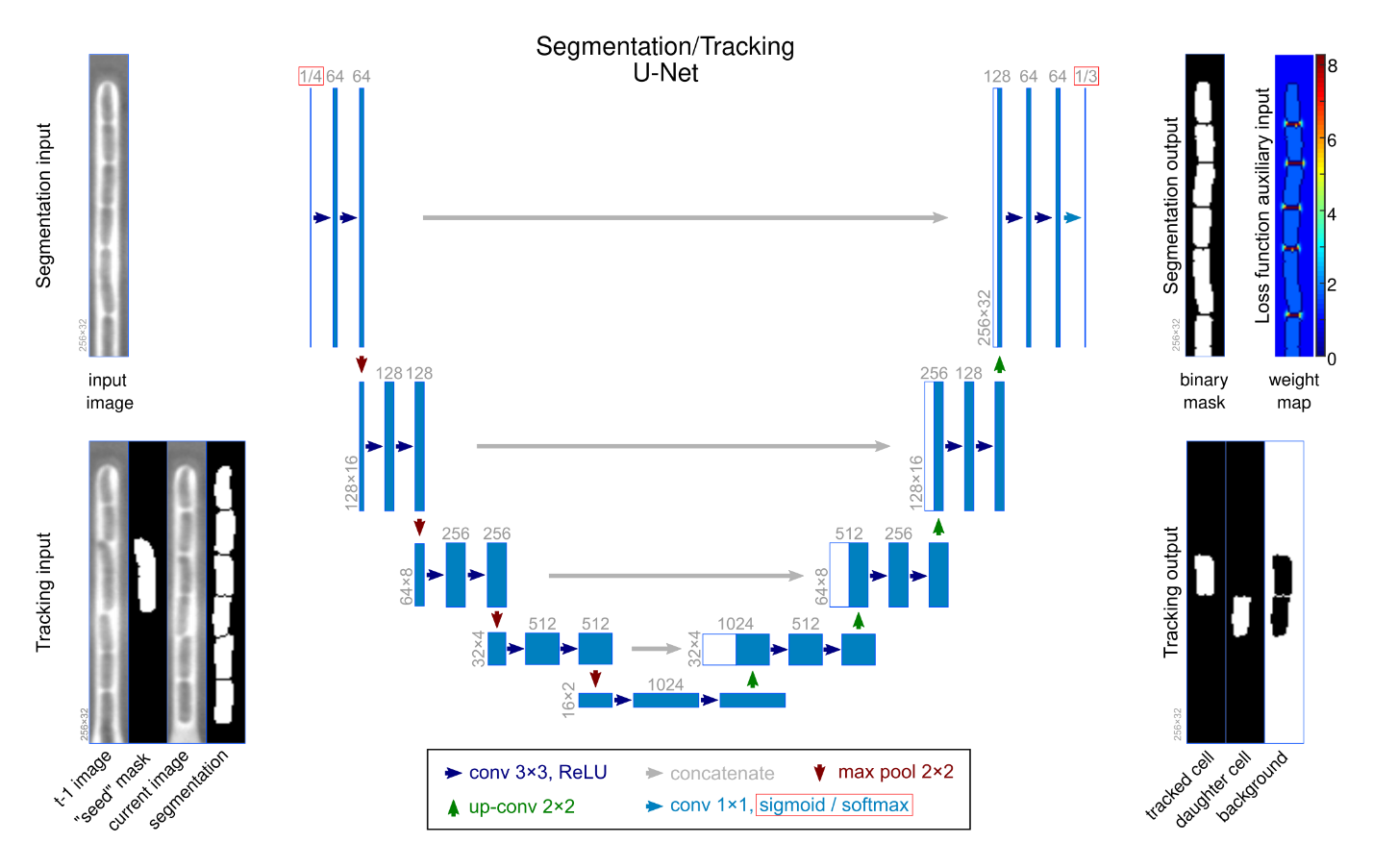

**Figure S5. U-Net architecture for segmentation/tracking.** The layers and tensor dimensions used in our U-Net implementation. The differences between the segmentation and the tracking U-Net are highlighted with red rectangular boxes. The first element is for the segmentation U-Net version, the second for the tracking version (segmentation/tracking). Note that the architecture is the same as the original U-Net model, except for image dimensions, number of input and output layers, and the final activation layer for tracking. The values we used for these are noted on the figure. The loss function for segmentation is a pixel-wise weighted binary cross-entropy. The pixel-wise weight maps are provided for each output mask during training as an auxiliary input. The loss function for tracking is a categorical cross-entropy.
